# Genetic control and tissue specificity of transcriptomic rhythmicity

**DOI:** 10.64898/2026.10.22.761195

**Authors:** Solal Chauquet, Valentin Hivert, Benjamin Weger, Frédéric Gachon, Loic Yengo, Sonia Shah, Allan F. McRae

## Abstract

Circadian rhythms influence molecular traits in most tissues, yet genetic studies mostly treat molecular measurements as static. In particular, genetic effects on gene expression are typically investigated through expression quantitative trait locus (eQTL) studies, which ignore sampling time and therefore miss genetic effects on rhythm parameters such as amplitude and acrophase. In this study, we used a cosinor-based joint genotype-by-rhythm interaction framework applied to GTEx version 10 gene expression data, with donor inferred phase as a proxy for circadian time. We first identified rhythmically expressed genes in 49 of the 50 tissues profiled in GTEx. We then tested common variants within 50 kb of each rhythmic gene’s transcription start site for genetic regulation of transcriptomic rhythmicity. We identified 167 circadian expression QTLs (cd-eQTLs), comprising 54 lead signals for 51 genes across 28 tissues. Most cd-eQTLs altered oscillation amplitude, while others affected acrophase or both. We then investigated the tissue specificity of cd-eQTLs and found that those acting on core circadian genes had consistent effects across tissues, whereas those acting on clock-regulated genes were largely tissue-specific. A phenome-wide association study in the Pan-UK Biobank linked 34 cd-eQTLs to 100 phenotypes, including self-reported chronotype, adiposity, blood cell counts and blood biochemistry. These results show that genetic variation shapes circadian gene expression in a tissue-dependent manner, and that accounting for time of day reveals regulatory effects that static eQTL analyses cannot detect.

## Introduction

A substantial fraction of the transcriptome exhibits circadian oscillation^1,2^; transcriptome-wide analysis estimates that roughly 43% of protein-coding genes are rhythmic in at least one tissue^3^. This rhythmicity is highly tissue-specific, with the liver displaying among the highest proportions of rhythmic transcripts (approximately 15% of its transcriptome^4^), while tissues such as the brain show considerably fewer rhythmic genes^3,5^.

At the cellular level, circadian rhythmicity arises from interconnected transcriptional-translational feedback loops (TTFL) that operate in most tissues^6,7^. The core loop is driven by the CLOCK:BMAL1 heterodimer, which activates transcription of the *Period* (*PER1, PER2*) and *Cryptochrome* (*CRY1, CRY2*) genes; the resulting PER:CRY protein complex then translocates to the nucleus and represses CLOCK:BMAL1 activity, generating a ~24-hour oscillation pattern^7^. A secondary stabilising loop, in which ROR and REV-ERB proteins compete to activate and repress *BMAL1* (*ARNTL*) and *CLOCK* transcription respectively, reinforces the amplitude and robustness of this rhythm^7^. Genes that constitute these feedback loops are named core circadian genes (CCG), while genes whose transcription is driven by CCG binding to E-box (BMAL1), RORE (REV-ERBs) or D-box (driven by clock-controlled PARbZip transcription factors) DNA motifs and related regulatory elements are termed clock-regulated genes (CRG)^8,9^. These intrinsic molecular clocks are present in almost all cells of the organism and are synchronised by a central pacemaker in the suprachiasmatic nucleus (SCN) of the hypothalamus and by environmental cues: the central clock in the SCN is synchronized by light^10^ while peripheral clocks are entrained by feeding^11^ and/or exercise^12^, ensuring alignment of internal biology with the external day-night cycle. In addition to environmental influences, genetic variation contributes to inter-individual differences in circadian timing, as reflected by heritable variation in chronotype^13^.

Beyond inter-individual variation in timing, disruption of circadian rhythms is associated with a broad range of health consequences. Misalignment between internal clocks and external schedules has been linked to increased risk of metabolic disorders^14,15^, cardiovascular diseases^14,16^ and cancer^17–19^. Psychiatric and behavioural traits, such as depression, bipolar disorder and neuroticism, have also been linked to circadian dysregulation^20^, with disrupted rhythmicity of core clock genes documented in major depressive disorder^21^.

However, despite this central role, most genetic studies of molecular traits capture only steady-state differences across individuals rather than their dynamic variation over time. For Cexample, expression quantitative trait locus (eQTL) studies^22,23^ typically do not account for the time of sample collection, treating individuals sampled at different points of the day as equivalent and thereby masking genetic effects that may act on rhythmic parameters such as amplitude (maximum value reached) or acrophase (time when the maximum value is reached).

To our knowledge, genetic effects on transcriptomic rhythmicity have been investigated by a single research group so far (Chen et al.^24^, Zhou et al^25^). Using the Genotype-Tissue Expression (GTEx) dataset version 8, Chen et al.^24^ assessed gene expression rhythmicity separately within each genotype group of a variant. They retained a variant-gene pair when rhythmicity was observed in at least one group (with p-value < 5 × 10^−4^ and peak-to-trough change > 1.5). Differential rhythmicity across genotypes was then tested with a joint test (HANOVA) on this subset of variant-gene pairs only. This test was further restricted to the two most frequent genotype classes, with a downsampling of the larger to the smaller observed sample size. Because the same data are used both to select variants and to test them, this sequential structure constitutes a form of selective inference bias^26,27^. Indeed, pairs are expected to be preferentially retained when sampling noise inflates the amplitude estimated in one genotype class, consequently inflating the apparent between-groups difference. This bias is amplified by stratification and downsampling, with smaller sub-groups yielding noisier amplitude estimates. Therefore, we expect this procedure to produce an excess of false positives.

In this study, we addressed these limitations and aim to systematically characterise genetic effects on rhythmic gene expression. For this, we used a joint interaction framework, analogous to the approach proposed in core chronobiological work^28^ alongside GTEx version 10^22,23^ gene expression and donor inferred phase (DIP)^5^ as a proxy for circadian time (Fig. 1a). This approach corrects for the selective inference bias exhibited in previously published work by jointly modelling the data. We identified rhythmically expressed genes across 50 tissues and tested the effect of nearby variants on rhythmic parameters, identifying loci we termed circadian expression QTLs (cd-eQTL). We then characterised the tissue specificity of cd-eQTL effects across CCG and CRG, and examined their phenotypic relevance through a phenome-wide association study in the Pan-UK Biobank^29^.

**Figure 1:**
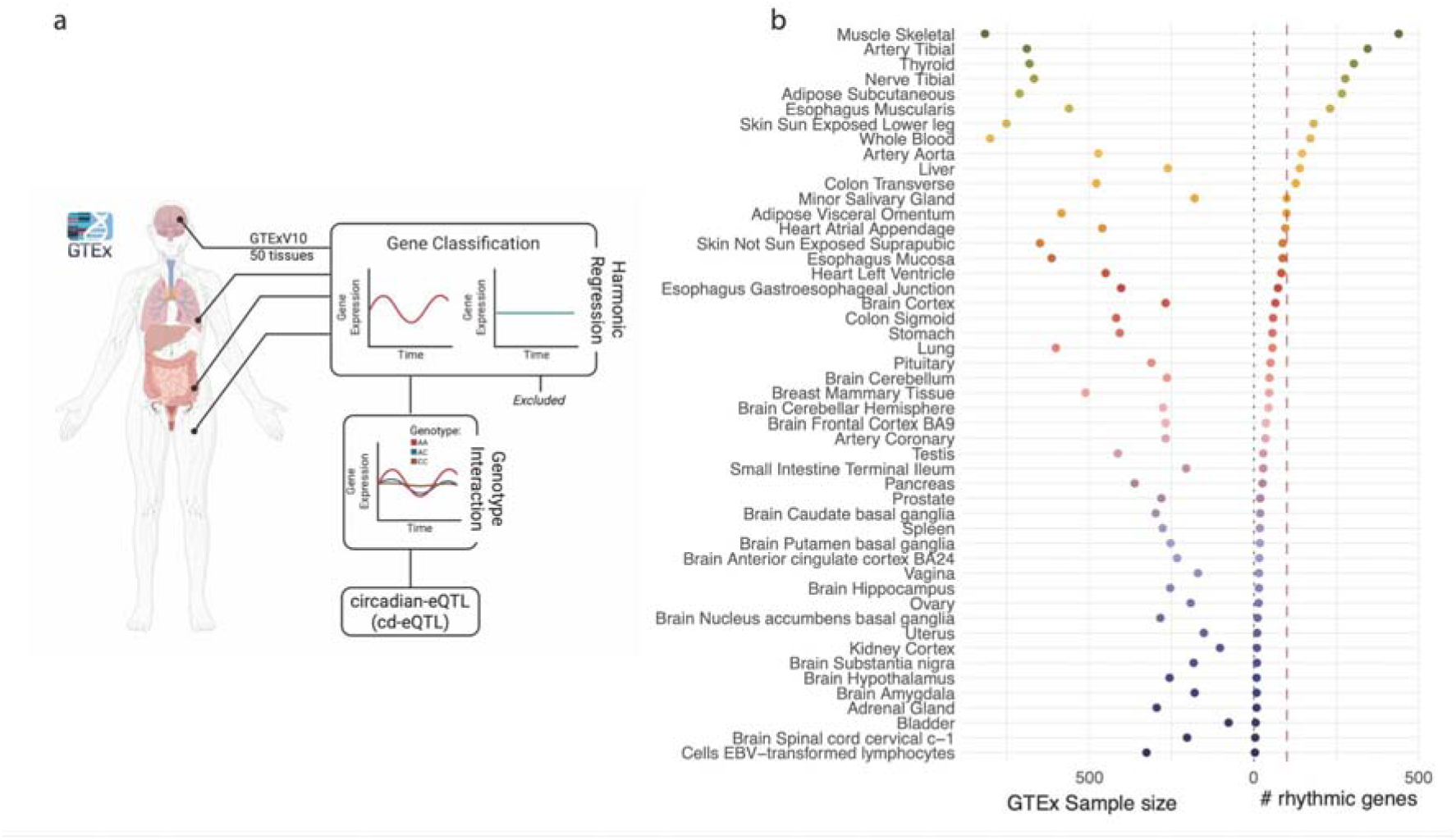
Identification of rhythmic genes in GTExV10. (a) Graphical abstract. GTExV10 was used to identify rhythmic genes by comparing a cosinor model to a constant model. The effect of all variants in a 100kbp window on rhythmic genes was then investigated using a cosinor model with an interaction term between rhythmic parameters and the variant of interest. All covariates defined by the GTEx consortium^62^ were used in all regressions. (b) Dotplot representing on the left the sample size in GTExV10 and the number of rhythmic genes identified in the same tissue on the right.

## Results

### Rhythmic genes

We started by identifying rhythmic genes within 50 different tissues from the GTExV10 dataset^22^. Rhythmicity was assessed by modelling gene expression as a function of DIP using a harmonic model for each gene within each tissue (*see Methods*). We then assessed goodness of fit using a log-likelihood ratio test (LRT) comparing the cosinor rhythmic model to a linear model, with both models including the same covariates (sex, sequencing platform, sequencing library protocol, 10 PEER factors and 5 genetic PCs). Genes whose harmonic model displayed a significantly better fit (LRT p-value < 0.05 after Bonferroni correction for the number of genes tested within each tissue) were considered rhythmic and retained for downstream genetic analysis. We identified at least one rhythmic gene in 49 tissues, with 11 tissues displaying more than 100 rhythmic genes (Fig. 1b, Supplementary Table 1). Skeletal muscle displayed the highest number of rhythmically expressed genes (439 significant genes), consistent with its well-documented rhythmicity^12^ as well as its larger sample size within GTExV10. Cultured fibroblasts were the only tissue without rhythmic genes, as expected. Indeed, cultured cells only exhibit rhythmicity once synchronised by an external stimulus such as serum^30^ or temperature^31^ shock, something that was not performed in the GTEx dataset.

### Genetic contribution to transcriptomic rhythmicity

Having identified rhythmic genes across tissues, we modelled the genetic contribution to rhythmicity by introducing an interaction term between genotype and cosinor parameters (Fig. 1a). For this, we selected a 100kbp window centred on each gene transcription start site (TSS) and tested the association between gene rhythmicity and all SNPs in this window. To account for the interaction term, which requires sufficient observations in each genotype class, we restricted the analysis to variants with minor allele frequency greater than 10%. For each variant-gene-tissue pair, we performed three regression analyses: one including the variant and covariates only, a second additionally including the cosinor parameters, and a third including the interaction between the variant and cosinor parameters. Goodness of fit was evaluated at each step using a LRT. We defined a significant cd-eQTL as one where the LRT p-value associated with the interaction model was smaller than 1 × 10^−4^, a threshold selected based on the expected number of independent linkage disequilibrium blocks in a 100kbp window^32^. Using this approach, we identified 167 cd-eQTL associated with 51 genes in 28 tissues. Retaining, for each gene in each tissue, a single variant with the lowest interaction p-value yielded 54 lead cd-eQTL (Fig. 2, Supplementary Table 2-4). The majority of tissues tested displayed one (n = 15) or multiple (n = 13) significant cd-eQTL associations, highlighting genetic control of circadian rhythmicity across several tissues. The tissue with the most cd-eQTL identified was Esophagus Muscularis (n = 7), followed by Tibial artery (n = 5). The associations fell into two distinct categories: CCGs (red highlight Fig. 2) and CRGs, with divergent patterns of tissue specificity. CRG associations were largely tissue-specific, while CCGs exhibited consistent genetic effects across multiple tissues. Next, we validated cd-eQTL identification using a recent model selection approach implemented in the dryR package^33^. Briefly, for 3 genotype groups, dryR fits 15 models (or 5 when the alternate homozygote group is observed in less than 10 individuals) ranging from one without oscillation to models where rhythmicity is either shared or independent between genotype groups. The model best representative of the data is then selected using a Bayesian information criterion weight, a probability like metric. Following this approach, 51 out of 54 (94%) independent cd-eQTL selected a model where rhythmicity was not shared between genotype groups validating the genetic contribution to rhythmicity (Supplementary Figure 1, Supplementary Table 5).

**Figure 2:**
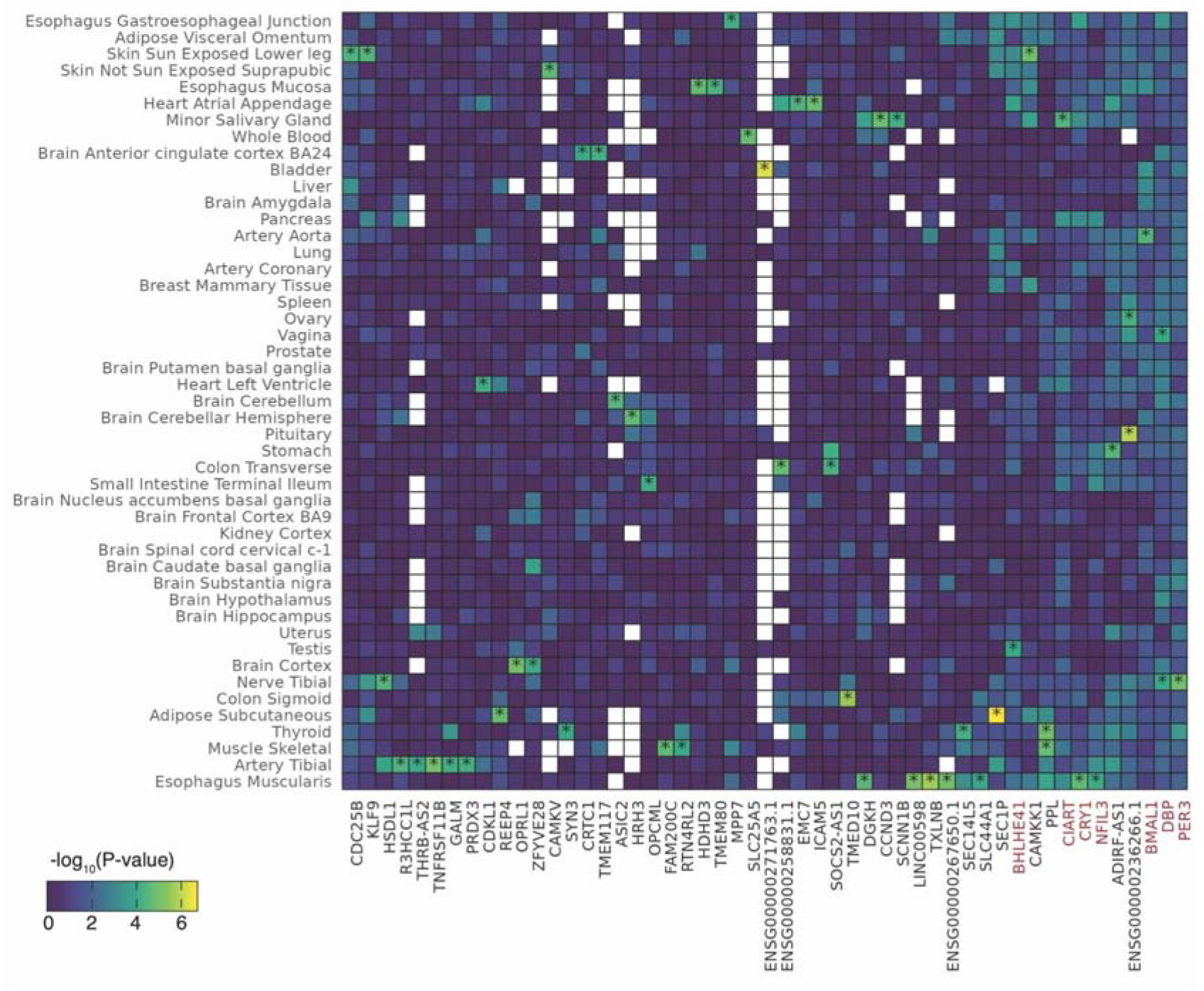
cd-eQTL can be identified in most tissues displaying rhythmicity. (a) Heatmap representing the −log_10_(p-value) of a log-likelihood ratio test comparing a model with rhythmic parameters and variant of interest, and a model including an interaction term between rhythmic parameters and variant. The p-value represents the significance of the effect of the variant on the estimated rhythmicity. Significance is set at p < 1 × 10^−4^ and is represented by a black star. A value of zero square represents a gene that did not display significant rhythmicity in the tissue of interest. White squares represent a gene not expressed in the tissue of interest. Gene names highlighted in red denote core circadian genes (CCGs).

We next characterised the nature of each genetic effect, asking whether a variant altered the amplitude of the oscillation, its acrophase (or time of peak expression), or both. The interaction model summarises the genetic effect as a single change in the sine and cosine coefficients; amplitude and acrophase are non-linear functions of these coefficients, so the model does not directly test whether they differ between genotypes, only that the overall rhythmicity change between genotype groups. We therefore fitted a separate cosinor to each genotype group on covariate-residualised expression, which estimates the amplitude and the acrophase of each group and compared them across genotypes using the phase and amplitude tests defined in Bingham et al,.^34^ To confirm that this residualised per-genotype cosinor captured the same genetic effect as the interaction model, we compared the amplitude and the acrophase estimated by the two approaches. The two estimates were highly concordant (Pearson r = 0.95 for amplitude and r = 0.92 for acrophase, Supplementary Figure 2). We then classified each cd-eQTL, assigning it to amplitude, acrophase, or both according to which test passed a false discovery rate of 0.1 across the 54 loci. As an acrophase is only defined for a rhythmic group, a phase effect further required at least two genotype groups with an amplitude different from zero. Three variants (rs7311890 on *TMEM117*, rs446246 on *SLC25A5* and rs142344547 on *CAMKV*) did not meet this requirement: only their largest genotype group had a detectable rhythm, so they could not be safely attributed to either an amplitude or a phase effect. Out of the 51 cd-eQTL where rhythmic parameters could be classified, amplitude effects were the most frequent: with 28 variants acting on the amplitude, 13 on the acrophase and 10 on both (Supplementary Table 6).

### cd-eQTLs influence both CCG and CRG

We identified a significant cd-eQTL influencing the rhythmic expression of *BMAL1*, a core component of the circadian transcriptional machinery, in the artery aorta (Fig. 3a). The variant rs11022718-T was associated with an allelic increase in rhythm amplitude, with each copy of the alternate allele corresponding to an ~0.25-unit increase (Fig. 3c). In contrast, the acrophase shifted modestly, from 21:09 in homozygous reference individuals to 23:47 in homozygous alternate individuals (Fig. 3a dotted line, Fig 3e). Similar genetic controls were observed for CRGs, with the expression of *TMED10* in the sigmoid colon modulated by rs76379942-Del (Fig. 3b). Here, the amplitude was unchanged in heterozygotes and increased by 0.22 units in homozygous alternate individuals (Fig. 3d). However, the predominant effect of this variant was on acrophase, with peak expression advancing from 22:39 in homozygous reference individuals to 19:13 in heterozygotes and 13:10 in homozygous alternate individuals (Fig. 3b dotted line, Fig. 3f). Together, these findings demonstrate that genetic variants can influence both amplitude and acrophase in both CCGs and CRGs.

**Figure 3:**
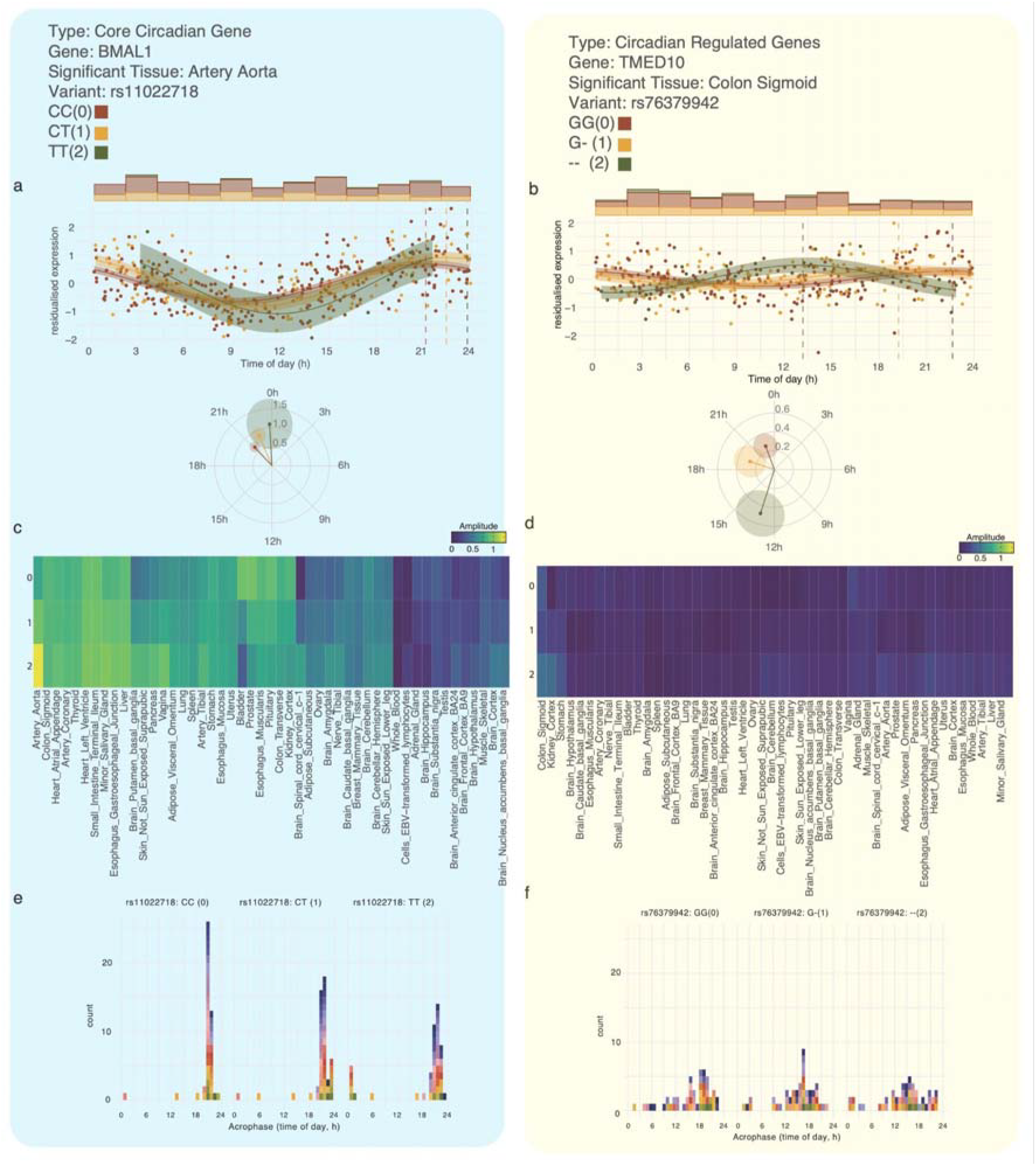
Example of a CCG and CRG displaying different behaviour across tissues. Rhythmicity of two significant cd-eQTL: *BMAL1*-rs11022718-T (left), a core circadian gene, and *TMED10*-rs76379942-Del (right), a clock-regulated gene. (a-b) Dot plot and polar plot showing the residualised gene expression of *BMAL1* and *TMED10* across genotype groups surrounded by its 95% confidence region. Dotted line represents the time at which expression peaks for a genotype group. Amplitude (c-d) or acrophase (e-f) calculated within genotype groups across different tissues for their respective genes. Each square corresponds to the acrophase of a tissue, the count corresponds to the number of tissue with a similar acrophase.

### Tissue specificity of cd-eQTL

Next, we investigated whether the observed genetic effects were tissue-specific or absent in other tissues due to a lack of statistical power. We modelled the expression of both *BMAL1* and *TMED10* across all 49 tissues that previously displayed at least one rhythmic gene. We observed a shared genetic regulatory effect of rs11022718-T on *BMAL1* in most tissues (Fig. 3c, e; Supplementary Tables 7, 8), while rs76379942-Del influenced *TMED10* in only a few tissues (Fig. 3d, f; Supplementary Tables 9, 10).

Seeing that CCG and CRG display different tissue specificity patterns, we conducted a systematic analysis of tissue specificity for both gene types. Using harmonic regression, we estimated four cosinor parameters for all significant cd-eQTL (sine, cosine, and their interaction terms with the variant of interest) and calculated pairwise cosine similarities between tissues based on these estimates, where a value of 1 indicated highly similar effects and −1 indicated opposing effects. For example, the effect of rs11022718-T on *BMAL1* was largely conserved across tissues (Fig. 4a, Supplementary Table 11), whereas the effect of rs76379942-Del on *TMED10* showed highly tissue-specific effects (Fig. 4b, Supplementary Table 12). To quantify these patterns, we averaged the cosine similarity across tissues for each cd-eQTL and compared the resulting distributions between the 8 CCGs and the 46 CRGs. CCGs displayed higher cross-tissue conservation (mean ± s.d. = 0.71 ± 0.12) relative to CRGs (0.19 ± 0.19), a difference that was significant (Wilcoxon rank-sum test, W=12, p=5.0×10^-7). We tested the robustness of this effect by recomputing the cosine similarity using only the interaction terms between the variant of interest and the rhythmic (sine and cosine) terms (Supplementary Figure 3). We observed a similar but weaker effect, bordering on significance (CCGs mean±SD = 0.15 ± 0.08; CRGs = 0.09±0.11; Wilcoxon rank-sum test, W = 106, p = 0.06). We argue that this lack of significance reflects limited statistical power, and that replication in a larger sample is warranted. Regardless, taken together these results suggest two distinct modes of genetic regulation of circadian rhythmicity: CCGs with consistent cross-tissue effects and CRGs with tissue-specific regulation.

**Figure 4:**
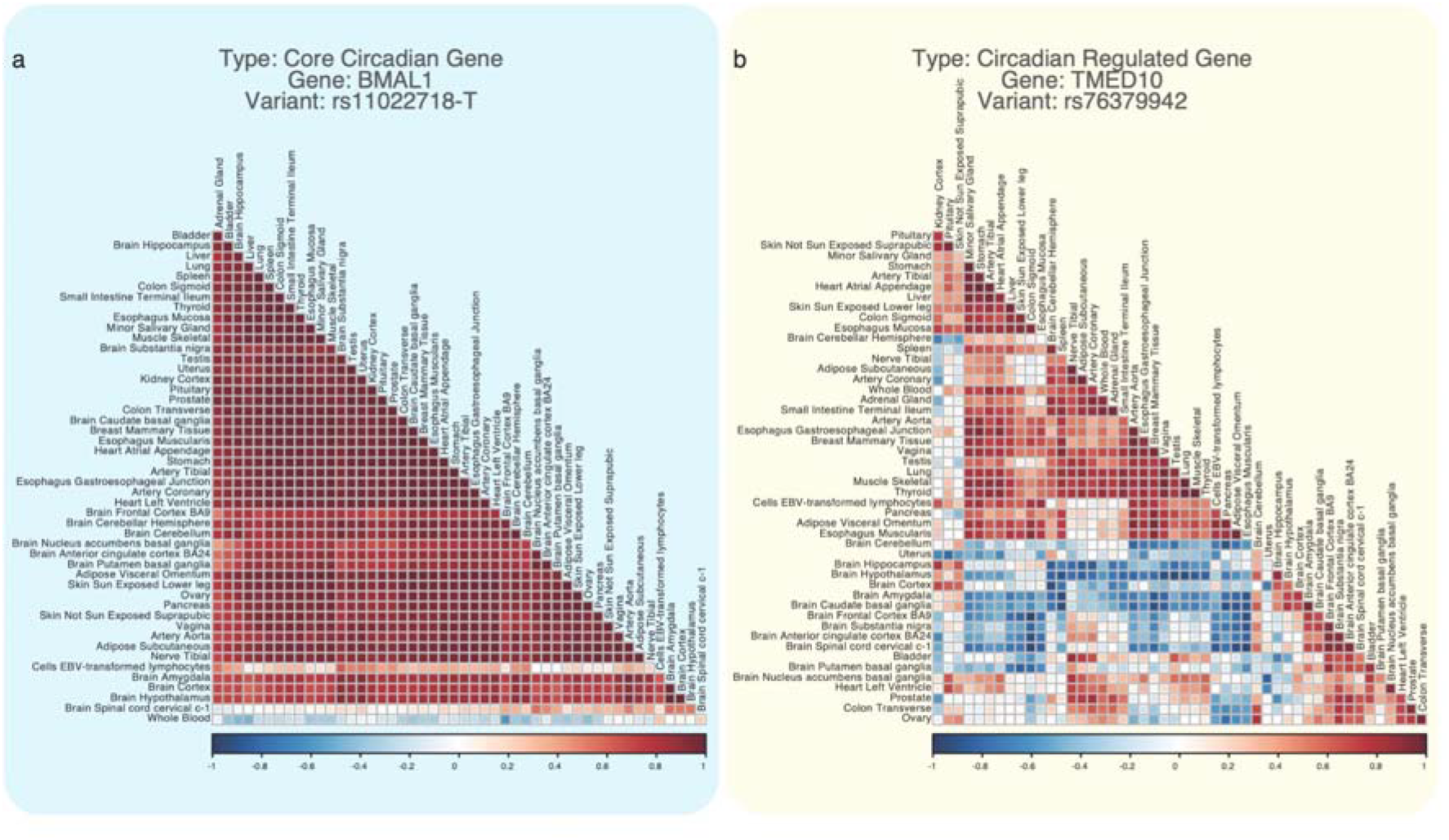
Tissue similarity calculated for a CCG and CRG. Cosine similarity of all four estimated rhythmic parameters representing the similarity of the effect of rs11022718-T on *BMAL1* (left) or rs76379942-Del on *TMED10* (right). A value of 1 represents a similar effect, −1 represents an opposite effect, and 0 an orthogonal one.

### Phenome-wide association study

Finally, we examined the biological relevance of cd-eQTL using a PheWAS approach. Of the 54 identified cd-eQTL, 49 were present in the Pan-UK Biobank GWAS^29^ (Supplementary Table 13). We extracted their associations across 1,085 phenotypes and grouped each trait into 21 categories, including blood biochemistry, blood count, physical measures, lifestyle and environment, as well as 17 ICD10 classes. Using a significance threshold of 4.6 × 10^−5^, we identified 310 associations covering 100 distinct phenotypes associated with 34 out of the 49 cd-eQTL.

Nearly half of these associations (49%) involved physical measures, while blood count and blood biochemistry accounted for 24% and 15% of associations, respectively. Additional signals were detected in lifestyle and environment traits: four cd-eQTL were associated with self-reported chronotype (strongest at rs697676-A, p = 2.95 × 10^−5^), three of which were also associated with ease of getting up in the morning, including the clock genes *PER3* and *CIART*. As an illustration, *BMAL1*-rs11022718-T (Fig. 5) was predominantly associated with physical measures of adiposity and body size (trunk fat percentage, p = 3.02 × 10^−3^; weight, p = 2.40 × 10^−^; hip circumference, p = 6.92 × 10^−^; body mass index, p = 9.51 × 10^−^), as well as with blood biochemistry (sex hormone-binding globulin, p = 1.68 × 10^−6^; total bilirubin, p = 1.62 × 10^−5^) and lifestyle and environment (plays computer games, p = 2.90 × 10^−6^).

**Figure 5:**
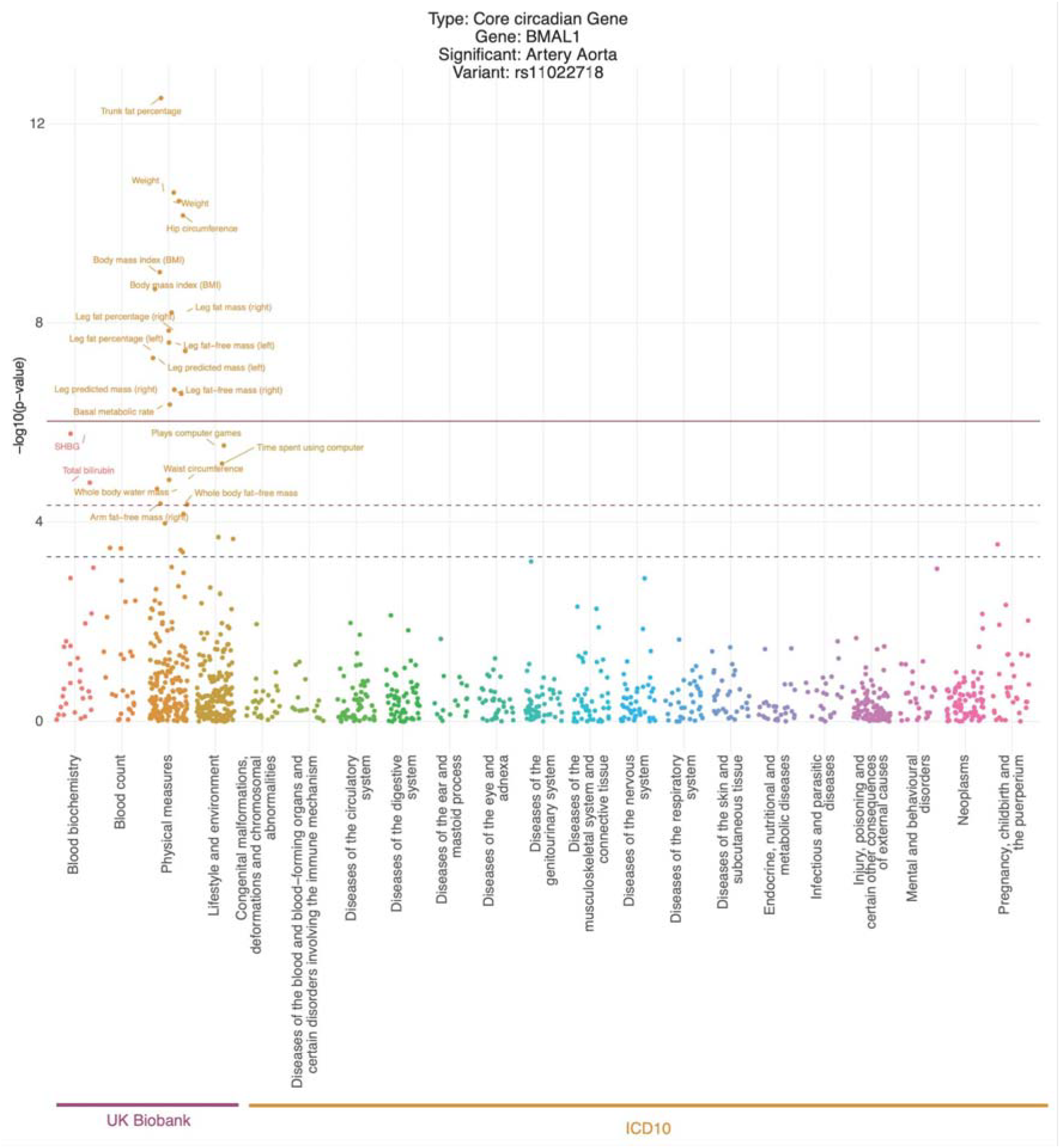
PheWAS of rs11022718-T, a cd-eQTL associated with *BMAL1* identified in Artery Aorta. Dotplot representing a PheWAS performed using 1,085 Pan-UK Biobank GWAS^29^. Phenotypes were sorted into 21 categories, including blood biochemistry, blood count, physical measures, lifestyle and environment, as well as 17 ICD10 classes. The dotted red line corresponds to the PheWAS significance threshold of 4.6 × 10^−^ while the dotted line corresponds to the suggestive threshold of 5 × 10^−4^ and a solid line representing the threshold when accounting for different cd-eQTLs tested of 8.53 × 10^−7^.

### Possible link between circadian clock gene rhythmic expression and chronotype

As described, two of the four cd-eQTL associated with chronotype, both of which were also associated with ease of getting up in the morning, act on the clock genes CIART (amplitude; Minor Salivary Gland, p = 4.50 × 10^−5^; Fig. 6a; chronotype p = 1.07 × 10^−4^) and PER3 (phase and amplitude; Nerve Tibial, p = 1.30 × 10^−5^; Fig. 6b; chronotype p = 1.29 × 10^−^). Both variants lie outside coding sequence: the CIART cd-eQTL (rs2147324) is in the 5′ untranslated region of exon 1, and the PER3 cd-eQTL (rs3135419) is in an intron of CAMTA1, upstream of PER3. The third cd-eQTL associated with both traits, rs697676 (chronotype p = 2.95 × 10^−5^; getting up in the morning p = 6.44 × 10^−6^), acts on ENSG00000236266, a transcript antisense to PER3(Fig. 6c). It lies upstream of the PER3 transcription start site and also altered the rhythmic expression of PER3, although this effect did not reach the cd-eQTL threshold (p = 7.90 × 10^−3^; Fig. 6d).

**Figure 6:**
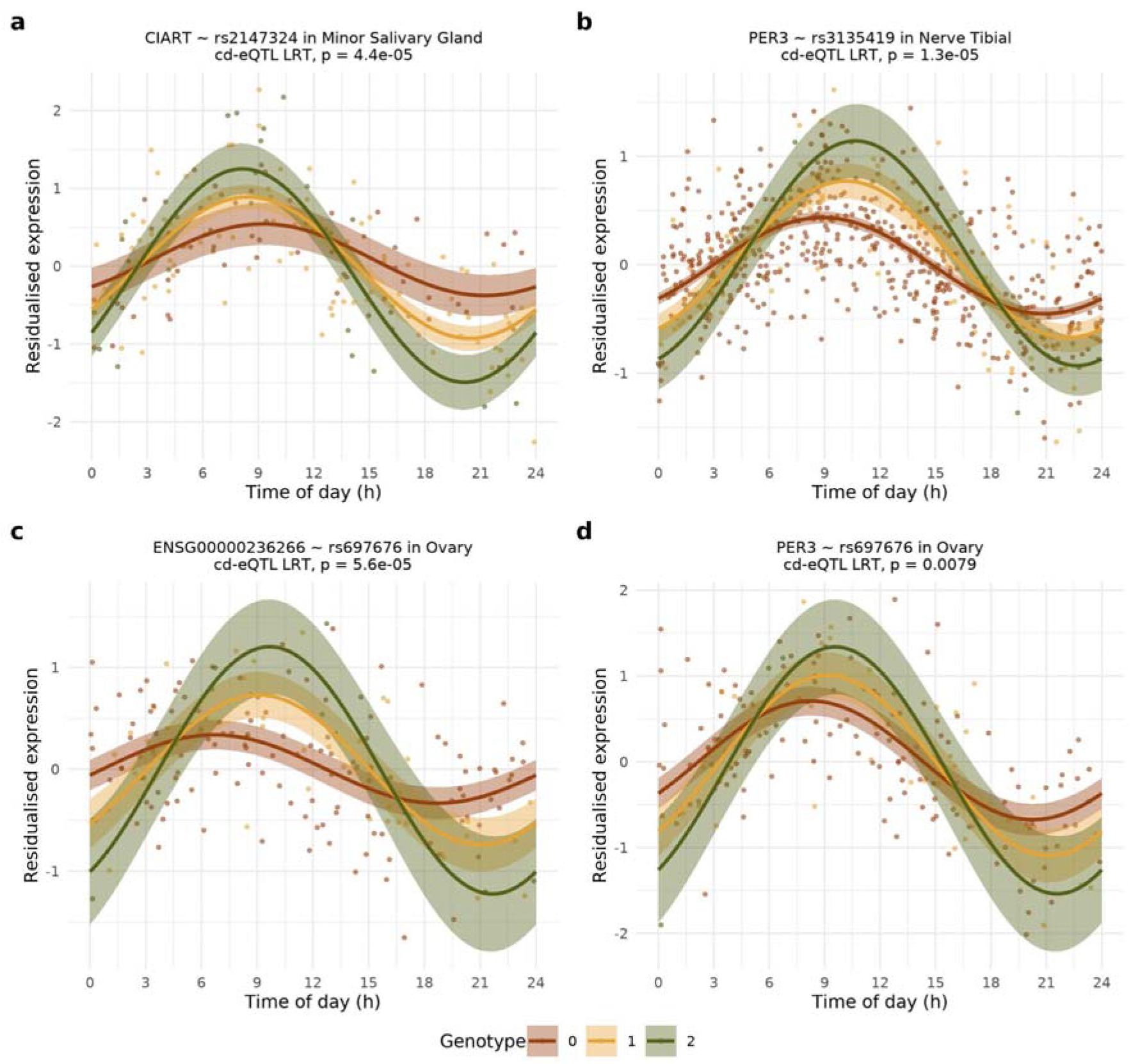
cd-eQTL of clock genes associated with chronotype. Dot plots of residualised gene expression against time of day by genotype group, with the fitted rhythm of each group and its 95% confidence band: (a) CIART-rs2147324-T in Minor Salivary Gland, (b) PER3-rs3135419-A in Nerve Tibial, (c) ENSG00000236266-rs697676-T, a transcript antisense to PER3, in Ovary, and (d) PER3-rs697676-T in Ovary, the tissue in which rs697676 is a cd-eQTL of ENSG00000236266.

## Discussion

Using DIP as a proxy for circadian time within the GTExV10 cohort, we detected rhythmic gene expression in 49 of 50 GTEx tissues, with skeletal muscle showing the largest number of rhythmic genes (439) and brain tissues showing only a few rhythmic genes. We then identified 54 lead cd-eQTL associated with 51 of these rhythmic genes across 28 tissues.

Chen et al.,^24^ reported several orders of magnitude more associations than we did: 3,027,405 significant variant-gene-tissue rhythmic QTLs (rhyQTLs), compared with our 167 cd-eQTL. At the tissue level, Chen et al. reported 183,758 rhythmic variants (6,790 independent loci) in visceral adipose tissue and 653 (210 loci) in substantia nigra, whereas we identified no cd-eQTL in either tissue. While this difference could be due to a different definition of the cis-window used (± 1 Mb compared to our ± 50 kb), restricting their associations to our window still retained 153,711 rhyQTLs. We argue that the remaining difference is largely due to selective inference bias^26,27^. Chen et al. tested differential rhythmicity only for pairs in which at least one genotype group was rhythmic, and only between the two most frequent genotype groups, after downsampling the larger group to the size of the smaller. Smaller subgroups yield noisier amplitude estimates, so pairs passing the filter carry larger noise-driven inflation. Low power in the remaining subgroup further encourages interpreting non-significant rhythms as absent, whereas a significant rhythm in one group and a non-significant rhythm in another does not establish a difference between them^27^.

To compare the two approaches directly, we selected the 4,513 rhyQTLs within our window that also passed our filters for population-level rhythmicity and allele frequency, and therefore had a matching test in our analysis (Supplementary Table 14). Only two of these rhyQTLs were cd-eQTLs in our analysis, *MPP7*-rs12220087 in the gastroesophageal junction) and *KLF9*-rs11142435 in sun-exposed skin, the latter being a SNP in high LD with our lead cd-eQTL variant rs11142441 (R^2^ = 0.976). For the remaining 4,511 rhyQTLs, our interaction model gave a median p-value of 0.31, only 13% reached nominal significance (p < 0.05), and our p-values were only weakly correlated with theirs (Spearman ρ = 0.11). Because our analysis used GTEx version 10 rather than version 8, the absence of most of our cd-eQTLs from their results could reflect the difference in sample size. We therefore re-implemented their pipeline on GTEx version 10 (Supplementary Table 15) and applied it to our 54 lead cd-eQTLs: only 25 (46%) were identified as significant, indicating that sample size alone does not explain the discrepancy. Finally, *AIF1* in heart left ventricle (Supplementary Figure 4), reported by Chen et al. as rhythmic only in rs7740525 GG carriers, showed a nominally significant rhythm in the GG group (p = 1.22 × 10^−3^, n = 143), but the genotype-by-rhythm interaction was not significant (p = 0.29). Taken together, these comparisons suggest that the joint interaction framework yields a smaller but more reliable set of cd-eQTLs than stratified approaches, which allows us to interrogate individual associations and their links to human phenotypes.

The cd-eQTLs associated with the two different types of circadian genes, CCGs and CRGs, displayed divergent patterns of tissue specificity, with CCGs showing conserved cross-tissue effects, while CRGs showed predominantly tissue-specific effects. This pattern is consistent with the known architecture of the mammalian clock: CCGs encode the core TTFL machinery active across most cell types^6,7^, variants affecting those are therefore more likely to produce conserved effects. CRG rhythmicity on the other hand, depends on tissue-specific co-regulatory programs^35^, restricting variant effects to the tissues where the relevant transcriptional context is present.

The *BMAL1* cd-eQTL, rs11022718-T, was observed to increase oscillatory amplitude by approximately 0.25 units per alternate allele while leaving the acrophase stable at around 21:00 across most tissues, highlighting the shared nature of cd-eQTL targeting CCG. *BMAL1* rhythmicity has been previously reported to be linked with psychiatric disorders^36^. Interestingly, rs11022718-T is nominally associated with neuroticism in a UK Biobank study^37^ (cross ancestry meta-analysis β = 0.044, SE = 0.0092, p = 1.86 × 10^−6^), and is in moderate linkage disequilibrium (D’ = 0.976, R^2^ = 0.356) with rs745752200-ATG, the top SNP associated with neuroticism (β = 0.055, p = 1.11 × 10^−11^) in the *BMAL1* locus. Additionally, rare variants found at the locus were associated with a delayed physical and neurological development through both amplitude-dampening and amplitude-increasing mechanisms^38^. The identified cd-eQTL could therefore play a mechanistic role in the effect of *BMAL1* in psychiatric disorders through a change in rhythmicity. Nevertheless, all these identified variants are localised in the coding sequence of the gene and rather impact its protein sequence. Here, we show that a genetic variant in a non-coding region and impacting *BMAL1* amplitude of rhythmic gene expression instead of its protein sequence can produce similar phenotypes in humans.

The *TMED10* cd-eQTL, rs76379942-Del identified in the sigmoid colon, illustrates the CRG pattern of tissue-specific effect. *TMED10* was previously identified as a rhythmic protein, supporting its behaviour as a clock-regulated gene^39^. The predominant effect of this variant was on acrophase: homozygous reference carriers peaked at 22:39 and homozygous alternate carriers peaked at 13:10, a shift of over nine hours. *TMED10* encodes a transmembrane p24 trafficking protein and plays a role in secretory pathways such as the unconventional protein secretion pathway^40,41^. Recent evidence showed that intestinal epithelial deletion of *TMED10* reduces IL-33 secretion, leading to severe experimental colitis^41^. Colitis is known to be linked to circadian disruption^42^, where reduced clock gene expression is observed in the inflamed intestinal mucosa of patients with inflammatory bowel disease^43^, and disruption of the colonic clock exacerbates experimental colitis in mice^44^. Additionally, in mice, circadian misalignment exacerbates experimental colitis and increases susceptibility to colitis-associated colorectal cancer^45,46^. *TMED10* rhythmicity may therefore be linked to the circadian biology of colitis, highlighting how cd-eQTL could shed light on disease mechanisms better than static eQTL.

We also examined the biological relevance of the cd-eQTL through a PheWAS in the Pan-UK Biobank, testing 49 of the 54 cd-eQTL against 1,085 phenotypes. As a positive control for our identification strategy, several cd-eQTL were associated with self-reported chronotype and ease of getting up in the morning. These chronotype associations are consistent with prior GWAS of morningness and sleep timing, which identified multiple loci near core clock genes^13^, and confirm that variants identified through rhythmic gene expression recover an established circadian behavioural phenotype. Having established this, we examined the broader phenotypic landscape. The PheWAS identified 310 associations covering 100 distinct phenotypes associated with 34 out of the 49 cd-eQTL. Those associations were linked to anthropometric and body composition traits (49%), blood count (24%), and blood biochemistry (15%) dominating. The *BMAL1*-rs11022718-T association illustrated how a single cd-eQTL can span multiple phenotypic domains, with associations found in adiposity and body size (trunk fat percentage, weight, hip circumference, BMI), sex hormone-binding globulin (SHBG) and total bilirubin. Several of these phenotypes are themselves linked to the circadian clock: *BMAL1* regulates adipogenesis and lipid metabolism^47^, and circadian disruption, such as shift work, is associated with obesity^48^, while circulating bilirubin varies over the 24-hour day^49^. In addition, in mouse models, *Bmal1* deletion leads to early obesity, altered fat storage, reduced body size, and alteration of sex hormones^50–52^. cd-eQTLs therefore map onto traits that are linked to circadian clock function and suggest that genetic variants impacting amplitudes can recapitulate some phenotypes observed in KO mouse models.

It is interesting to note that non-coding variants associating rhythmic amplitude with phenotypes are not specific for *BMAL1*. cd-eQTLs impacting *CIART* and *PER3* amplitude and/or phase have also been linked with chronotype and difficulty to “getting up in morning”. While several genetic variants of *PER2*^53^, *PER3*^54,55^, *CRY1*^56^, or *CRY2*^57^ have been associated with chronotype in humans, they all impact the sequence of the protein and its biochemical properties. Here we show that non-coding variants can also modify the chronotype by impacting the rhythmic expression of *CIART* and *PER3*. Because the phase of *PER3* expression directly correlates with the chronotype^58^, this suggests that genetic variants impacting *PER3* phase can have a direct role in modulating the chronotype. Accordingly, PER3 variants were identified as the most significant hits in several GWAS studies for chronotype^59,60^. Nevertheless, and according to our knowledge, the only non-coding mutation impacting the circadian period and phenotype is the deletion of an intronic enhancer in the *Cry*1 gene in mice^61^. Disrupting the rhythmic chromatin loop formed by the promoter and the enhancer leads to a decreased and phase-advanced *Cry1* expression and a shortening of the circadian period. Taken together, the PheWAS signals are consistent with known circadian effects and possibly hints at the tagging by GWAS studies of circadian variation in measurement rather than real effect of variants on traits. Moreover, this suggests that these variants could function as transcriptional enhancers and deserve to be functionally studied in the future.

Finally, our study has a few limitations. Firstly, our analysis relied on donor inferred phase as a proxy for circadian time, an estimated rather than a directly measured metric. Second, we restricted testing to genes rhythmic at the population level, to common variants (MAF > 0.10), and to variants within 50 kb of the TSS, owing to the statistical power requirements of the interaction framework. Genetic effects that confer rhythmicity on otherwise non-rhythmic genes, the contribution of rare variants, and distal or trans-acting effects therefore remain biologically relevant but outside the scope of this study. To our knowledge, no dataset currently samples the full 24-hour day at a scale sufficient to relax these constraints. Generating a circadian atlas would allow the effects of cd-eQTL to be resolved more finely, and would open the way to quantifying the circadian contribution to classical genetic metrics such as heritability.

## Acknowledgement

We wish to acknowledge The University of Queensland’s Research Computing Centre (RCC) for its support in this research

## Funding Statement

L.Y. is supported by the Snow Medical Research Foundation and the Australian Research Council (FT220100069). S.S. was supported by the National Heart Foundation Future Leader Fellowship (105638). F.G. is supported by the Novo Nordisk Foundation (Hallas-Møller Ascending Investigator grant 0087882).

## Methods

### GTEx data

RNA data stemming from 946 donors covering 50 different tissues were used for the analysis. Sample size distribution by tissue can be found in Figure 1b.

#### Genotype data

Whole genome sequencing (WGS) data was obtained from 946 individuals at a median depth of 32x. Samples were sequenced using either an Illumina HiSeq 2000 with a PCR-based protocol or an Illumina HiSeq X with a PCR-based or PCR-free protocol. WGS data were then processed with a pipeline described previously^62^. Genotype principal components (PCs) were previously computed using EIGENSTRAT^63^ and made publicly available (GTEx_Analysis_v10_eQTL_covariates.tar).

#### mRNA data

mRNA sequencing data were already formatted for QTL analysis by the GTEx consortium and were obtained from the GTExV10 release (GTEx_Analysis_v10_eQTL_expression_matrices.tar). Briefly, RNAseq data were aligned to the GRCh38 human genome with STAR^64^. Low quality samples were removed based on the metric reported previously^62^. Mapped reads were normalised between samples using a trimmed mean of M values (TMM) approach^65^. Genes passing the following criteria were then kept for analysis: ≥0.1 TPM in ≥20% of samples, and ≥6 reads (unnormalized) in ≥20% of samples. Finally, the expression values for each gene were inverse normal transformed across samples. The resulting normalised gene expression was then used to perform quantitative trait loci analyses.

#### Covariates

The Probabilistic Estimation of Expression Residuals (PEER) method^66^ was previously used by the GTEx consortium to account for hidden technical effects in the gene expression data. We used pre-computed PEER factors for the GTEx version 10 release (GTEx_Analysis_v10_eQTL_covariates.tar). In addition to latent factors, covariates previously demonstrated to be associated with gene expression^62^ were obtained from the GTEx version 10 release (GTEx_Analysis_v10_eQTL_covariates.tar). Namely, donor sex, whole genome sequencing platform (HiSeq 2000 or HiSeq X), whole genome sequencing library construction protocol (PCR-based, PCR-free), 10 tissue specific PEER factors and 5 genetic PCs were used as covariates in the QTL analysis.

### Proxy for circadian rhythmicity

Circadian analyses require a reliable proxy for circadian time. Although the GTEx dataset includes time of death (TOD), it is an imperfect circadian proxy since it does not account for heterogeneity in individual chronotype, time zone or cause of death^67^. To address these limitations, several methods have been developed to infer donor circadian phase directly from RNA sequencing data, including CHIRAL, which was specifically designed for the GTEx dataset^5^. As donor inferred phase (DIP) provides a more accurate proxy for circadian time, we used it throughout our analysis. DIP values were obtained from the published CHIRAL study^5^. Although DIP was inferred using GTEx release v8, whereas the present study uses GTEx release v10, the increase in sample size between these releases was driven primarily by additional RNA-seq samples from existing donors, with only 32 newly sequenced individuals added in v10.

Consequently, the published DIP estimates were used directly without re-estimating donor phases.

### Harmonic gene selection

To reduce the multiple testing burden and limit the number of false positives, we restricted our analyses to genes exhibiting evidence of rhythmic expression. Accordingly, we first identified rhythmic genes in the GTEx dataset and subsequently performed QTL mapping only for this subset of genes. To identify rhythmic genes, we fitted both a linear regression and a cosinor (harmonic) regression^34^ of the following form:

- 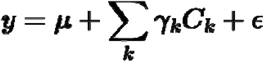
- 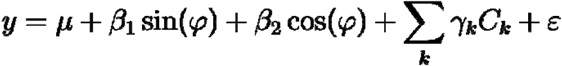

Where:

- ***y*** is the inverse-normal-transformed expression of the gene within a specific tissue.
- ***μ*** is the intercept, also called mesor in the cosinor models.***γ*_*k*_**
- ***C*_*k*_** are the p = 18 covariates with their associated coefficients
- ***φ*** corresponds to the radian-transformed DIP.
- ***ε*** is the residual error of the model.

The linear model assumes that gene expression remains constant over time, whereas the cosinor model represents gene expression as a sinusoidal function of circadian time. Model fit was evaluated using a likelihood ratio test implemented in the lmtest R package^68^ (lrtest function), with the null hypothesis that the linear model provides an adequate fit to the data. The resulting likelihood ratio statistics were compared with a chi-squared distribution with two degrees of freedom, corresponding to the difference in the number of model parameters between the two models. P-values were adjusted for multiple testing using a Bonferroni correction for the number of genes within each tissue. Genes for which the harmonic model provided a significantly better fit than the linear model (Bonferroni-adjusted p < 0.05) were retained for downstream QTL mapping. This procedure was performed independently for each tissue.

Importantly, our approach was designed to identify genes exhibiting rhythmic expression at the population level before testing for genetic effects on rhythmicity. As a result, we did not evaluate genes that are non-rhythmic in the overall population but become rhythmic only in specific genotype groups (i.e., conditionally rhythmic genes). Detecting such genotype-dependent rhythmicity would require directly testing genotype-by-time interactions across all genes, substantially increasing the multiple testing burden and reducing statistical power to detect these effects reliably.

### QTL regression

After identifying genes exhibiting significant harmonic expression patterns, we conducted QTL mapping for all SNP located within 50,000bp upstream and downstream of each gene transcription start site (TSS), corresponding to a 100kbp window centred on the TSS. Since harmonic regression requires uniform temporal coverage to reliably detect rhythmic patterns over the 24-hour cycle, we applied a more stringent minor allele frequency (MAF) threshold than typically used. Specifically, variants with MAF below 10% were excluded to ensure adequate representation of all genotype-time combinations.

We then identified significant circadian eQTLs by fitting three models for each gene-variant pair:

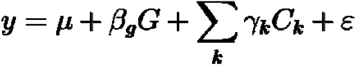

1. Linear model:

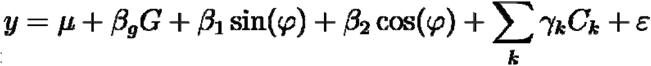
2. Harmonic model:
3. Harmonic interaction model:

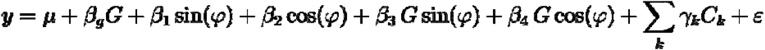

Where:

- **y, *μ,C_k_, φ*,** and ***ε*** are the same as previously defined.
- ***G* ∈ {0,1,2}**is the genotype of the variant tested, coded as the alternate-allele dosage, with main effect *β*_*g*_
- **β_3_, β_4_** are the coefficients of the genotype-by-rhythm interaction terms ***G* sin (*φ*)**and ***G* cos (*φ*)**, which capture the genetic effect on the rhythm and are absent from models 1 and 2.

The set of covariates was the same as used in the harmonic gene selection analysis. Model comparisons were performed using sequential likelihood ratio tests. Gene-variant pairs were classified as significant cd-eQTLs when the likelihood ratio test of the harmonic interaction model (model 3) against the harmonic model (model 2) yielded a p-value, indicating that the genotype-by-time interaction terms significantly improved the fit over the model containing the variant and rhythm without interaction. This threshold corresponds to a Bonferroni correction for the approximate number of independent variants within a 100 kb window^32^. To identify lead cd-eQTL signals, we conserved a single variant per tissue-gene combination with the most significant LRT p-value was retained.

### DryR validation

We validated cd-eQTL identification using a model selection approach implemented in the dryR package^33^. Briefly, for 3 genotype groups, dryR fits 15 models (or 5 when the alternate homozygote group is observed in less than 10 individuals) ranging from one without oscillation to models where rhythmicity is either shared or independent between genotype groups. The model best representative of the data is then selected using a Bayesian information criterion weight, a probability like metric. Following the recommendations from the paper, the model with the highest probability was selected as the model best representing the rhythmicity.

### Rhythmicity Parameter

Rhythmic gene expression can be fully characterised using two parameters:

- Amplitude: the maximum deviation of gene expression from its mean level (mesor) over the cycle
- Acrophase: the time at which gene expression reaches its peak

However, cosinor regressions do not directly estimate phase and amplitude. For a harmonic regression of the form:

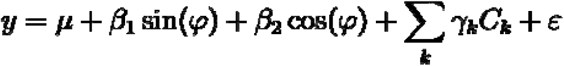

The parameters are recovered as:

- Amplitude: 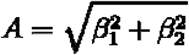
- Phase: **Φ = atan2(*β*_1_ *β*_2_)**

In the context of a model with interaction terms such as the one used to identify cd-eQTL:

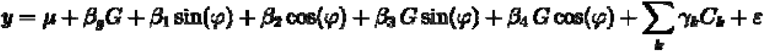

The rhythmicity parameters are estimated as a function of the inferred regression parameters and genotype group : ***G* ∈ {0,1,2}**

- Amplitude: 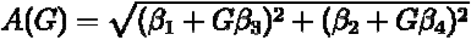
- Phase: **Φ (*G*) = atan2(β_1_ + *G*β_3_, β_2_ + *G* β_4_)**

### Confidence intervals for rhythmicity parameters

Amplitude and acrophase are non-linear functions of the fitted coefficients, so we derived their uncertainty following the inferential cosinor framework of Bingham et al^34^.

For each genotype group ***G***, the rhythm vector **b(*G*) = (*β*_1_ *+ Gβ*_3_, *β*_2_ + *G β*_4_)** has a covariance **∑(*G*)** obtained from the covariance of the rhythm coefficients**(*β*_1_,*β*_2_*β*_3_*β*_4_)**. From **∑(*G*)** we computed 95% confidence intervals for the amplitude and acrophase using the delta method, as well as their joint 95% confidence ellipse in the **(sin,cos)**coefficient plane, in which the distance from the origin represents the amplitude and the polar angle the acrophase.

### cd-eQTL classification

To classify each cd-eQTL according to whether the genotype influenced the amplitude, the acrophase, or both, we applied the cosinor parameter tests of Bingham et al^34^. Because the inferential framework does not allow for covariates, we first regressed gene expression on the full covariate set in a single model fitted across all samples and carried the residuals forward. We then fitted an independent cosinor model to each genotype group on these residuals, so that every group contributed its own amplitude and acrophase. We tested the equality of the amplitudes and, separately, of the acrophases across the genotype groups using the F-tests of Bingham et al., which provided one amplitude and one acrophase p-value per cd-eQTL. These p-values were adjusted for multiple testing across all cd-eQTL using the Benjamini-Hochberg procedure, and each variant was classified as acting on amplitude, acrophase, or both according to which adjusted p-values reached significance (FDR < 0.1).

### Tissue specificity estimation

To quantify the similarity between tissues, we performed a cosinor regression using the model with a genotype-by-time interaction for each of the identified cd-eQTL in all tissues. For a specific variant-gene combination, this approach generated a vector of 4 values corresponding to the estimated parameters for the sin, cos, sin-by-genotype and cos-by-genotype interactions. We then calculated the pairwise cosine similarity between all tissues. A value of 1 indicates that all estimated coefficients were similar between tissues, representing a shared genetic signal, while a value of −1 represents a fully opposite genetic effect. A value of 0 does not imply a lack of rhythmicity, but an orthogonal effect to the one observed in the compared tissue.

We then used the mean similarity value across all comparisons as a proxy for a shared signal between tissues. We grouped genes into CCG and CRG and compared the mean cosine similarity across cd-eQTL using a Wilcoxon rank-sum test.

### Phenome-wide association study

We performed the PheWAS analysis using data from the Pan UK Biobank study including 420,531 unrelated European individuals. While the Pan UK Biobank includes meta-analysis of different genetic ancestries, we limited our analysis to European ancestry for concordance with the GTEx dataset. The detailed quality control procedures for genotype data and selection of European individuals are described in Karczewski et al.^29^. Briefly, GWAS summary statistics for the variants identified as cd-eQTL were extracted for 1,085 phenotypes belonging to one of the following classes: Blood biochemistry, Blood count, Physical measures, Lifestyle and environment, and 17 ICD10 classes. Associations were considered significant when passing the conservative threshold of 4.6 × 10^−5^. This threshold was obtained by performing a Bonferroni correction based on the number of traits extracted, assuming that they were all independent.

### Comparison to rhyQTL

To test whether our cd-eQTL would be detected by the approach used by Chen et al., we re-implemented the rhyQTL pipeline from their publicly available code and applied it to the 54 lead cd-eQTL. For each variant, donors were grouped by genotype and only the two most frequent genotype groups were retained. Rhythmicity was assessed in each group by harmonic regression, requiring p < 5 × 10^−4^ and a peak-to-trough ratio > 1.5 in at least one group. Differential rhythmicity was then assessed with dryR after downsampling the larger group to the size of the smaller one using dryR; variants assigned to models with no rhythm or with a rhythm shared between groups were excluded. Finally, rhythm differences were tested with HANOVA (p < 0.05). Of note, we used residualised GTEx v10 expression rather than log2 CPM (Supplementary Table 15).

## Code availability

All code used to perform the analyses reported in this paper can be found on the GitHub following repository: https://github.com/SolalC/cd-eQTL

## Supplementary Figures

**Supplementary Figure 1.**
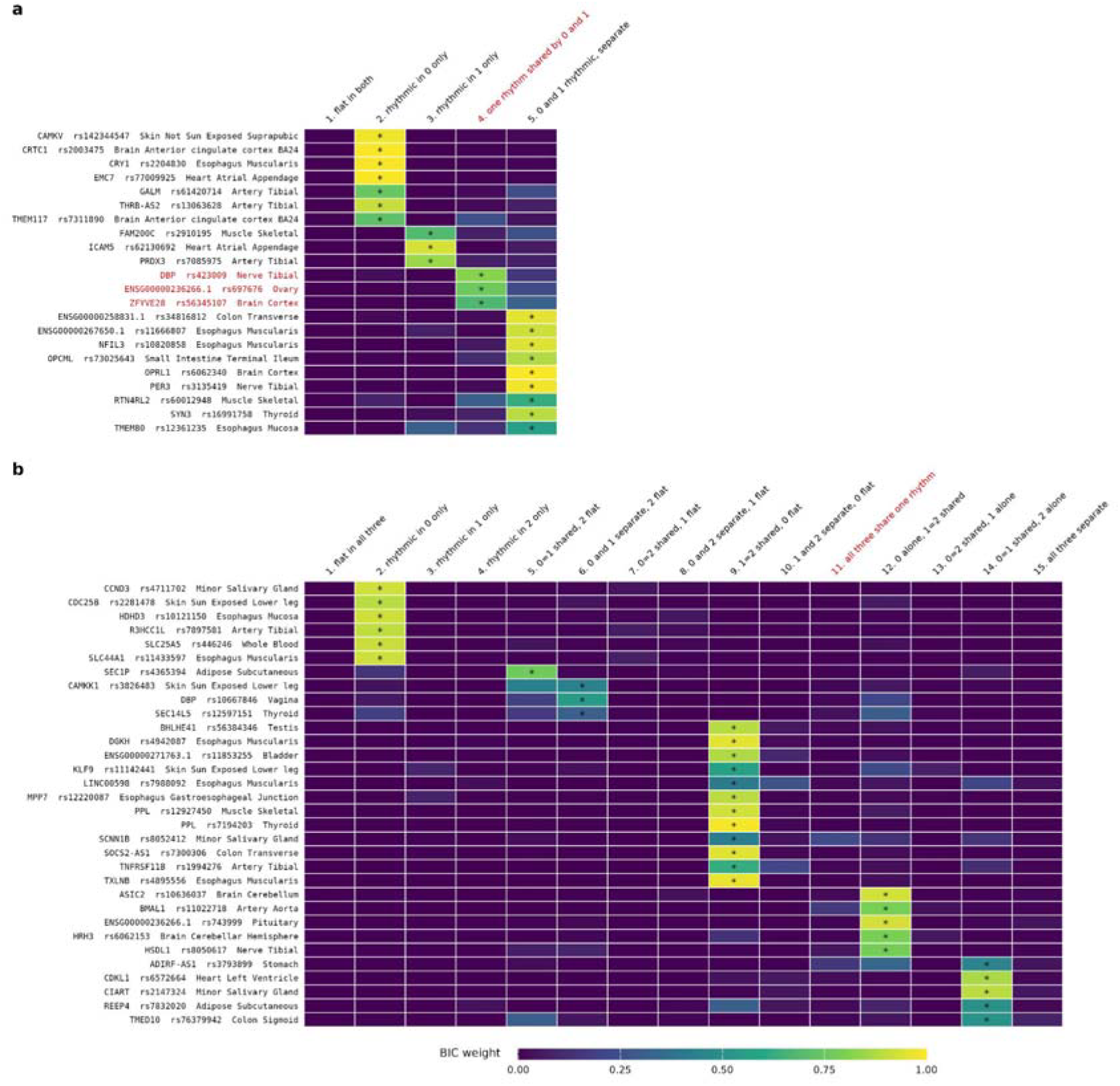
Rhythm model selection by dryR at the 54 lead cd-eQTL. (a) Loci with two usable genotype groups (0 and 1), scored against 5 candidate models. (b) Loci with three usable genotype groups (0, 1 and 2), scored against 15. Each row corresponds to a tissue-gene-variant combination; each column is one of dryR’s candidate rhythm models. Cell colour is that model’s BIC weight, which sums to 1 across the models of a locus, the star marks the model selected. Genotype groups were less than 10 donors where present were removed, leading to some comparison including only two genotype groups. The model and the cd-eQTL associated where all genotype groups share one rhythm (no genotype effect on rhythmicity) are labelled in red.

**Supplementary Figure 2:**
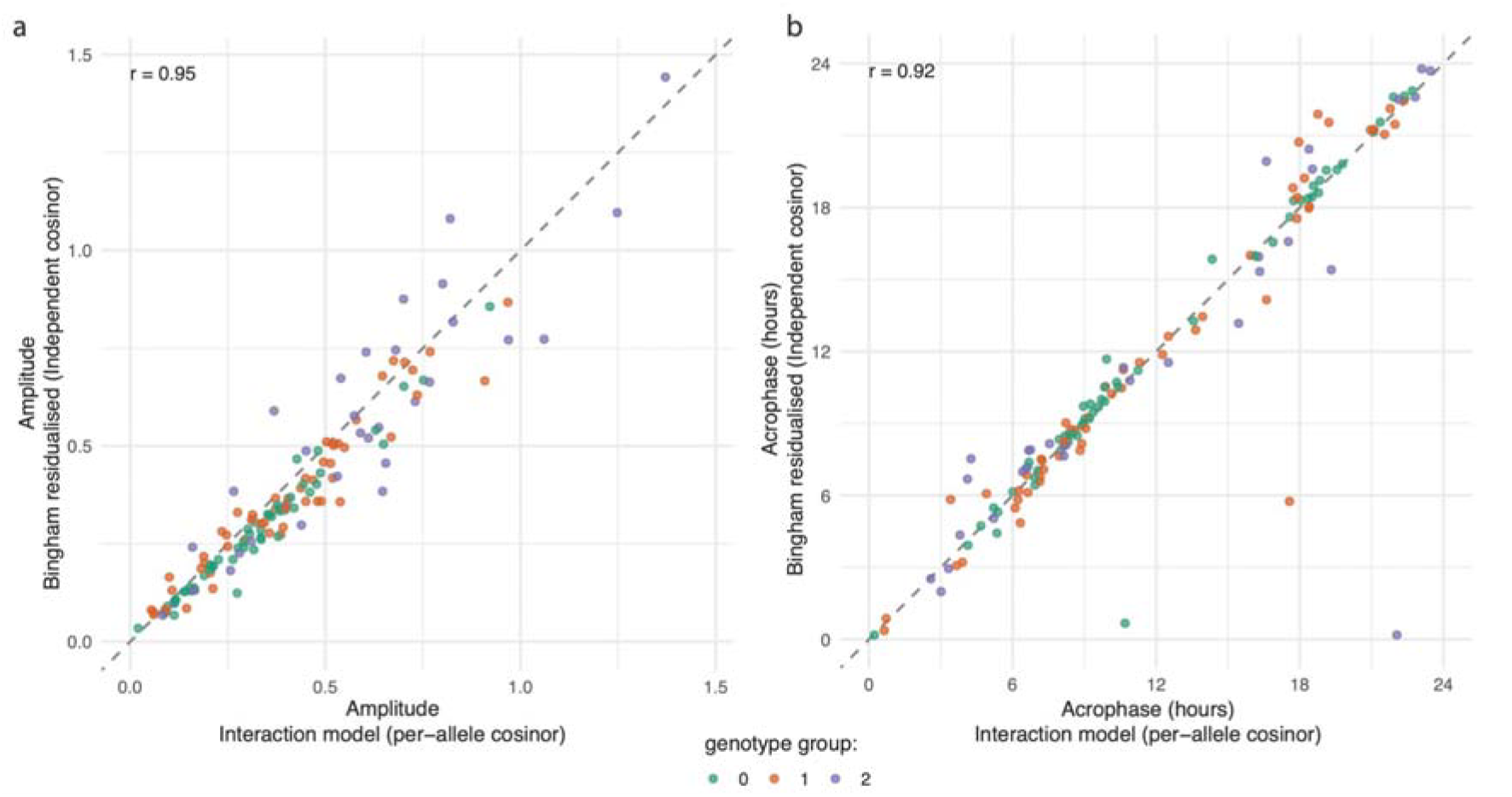
Comparison between independent and interaction rhythmicity parameters. For each genotype group of the 54 lead cd-eQTL, the amplitude (a) and the acrophase (b) estimated from the interaction cosinor approach used to detect cd-eQTL and the residualised per-genotype cosinor used to compare rhythmic parameters were compared. Points are colored by genotype groups. The dashed line represents the identity (y=x) line. The two estimates are highly concordant, highlighting that the classification based on residualised cosinor reflects the same genetic effect captured by the interaction model.

**Supplementary Figure 3.**
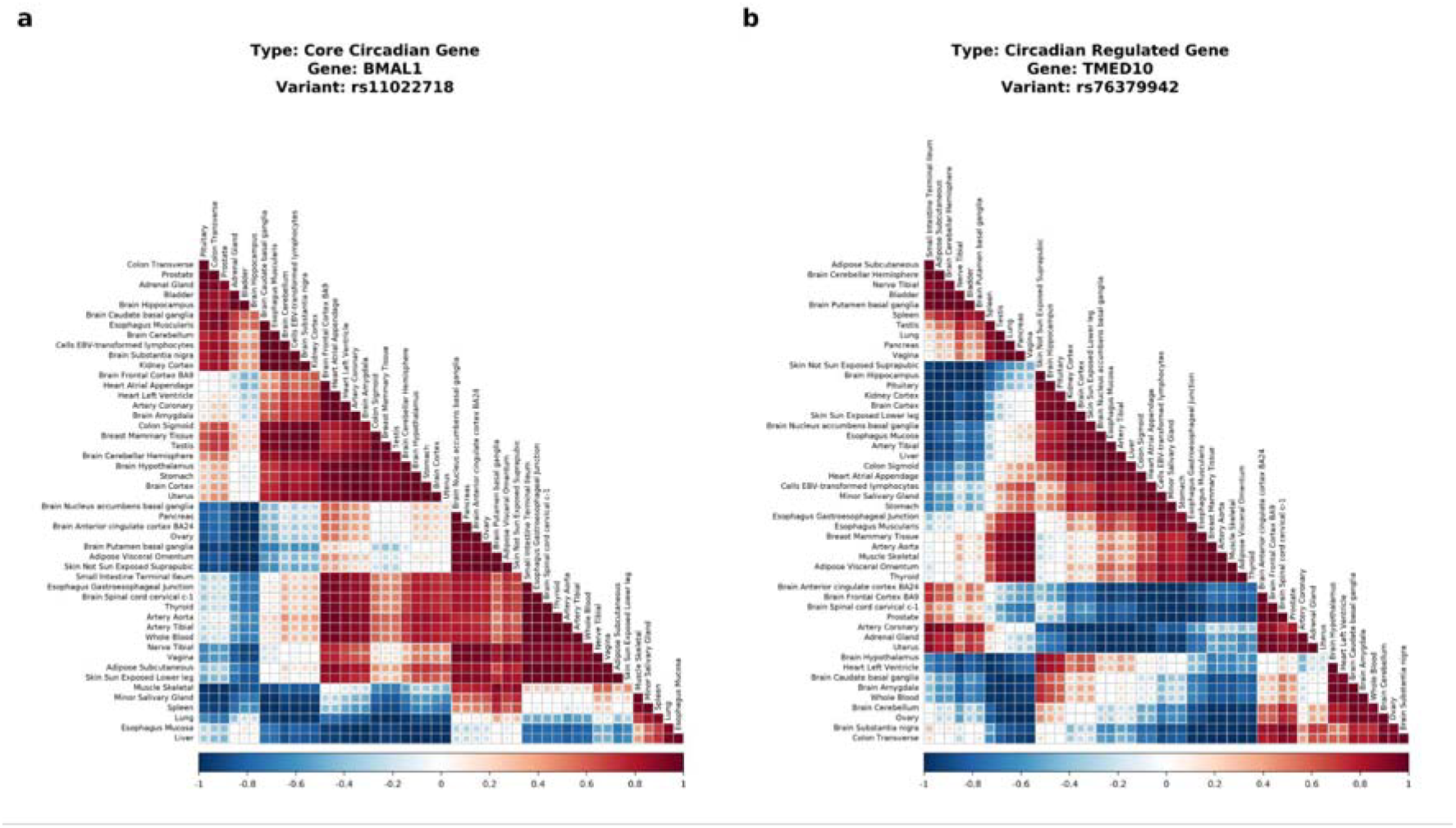
Cross-tissue cosine similarity of the cd-eQTL effect computed on the genotype-by-time interaction terms only. (a) rs11022718-T on *BMAL1* (mean pairwise similarity 0.23). (b) rs76379942-Del on *TMED10* (mean pairwise similarity 0.00). A value of 1 represents a similar effect, −1 represents an opposite effect, and 0 an orthogonal one.

**Supplementary Figure 4:**
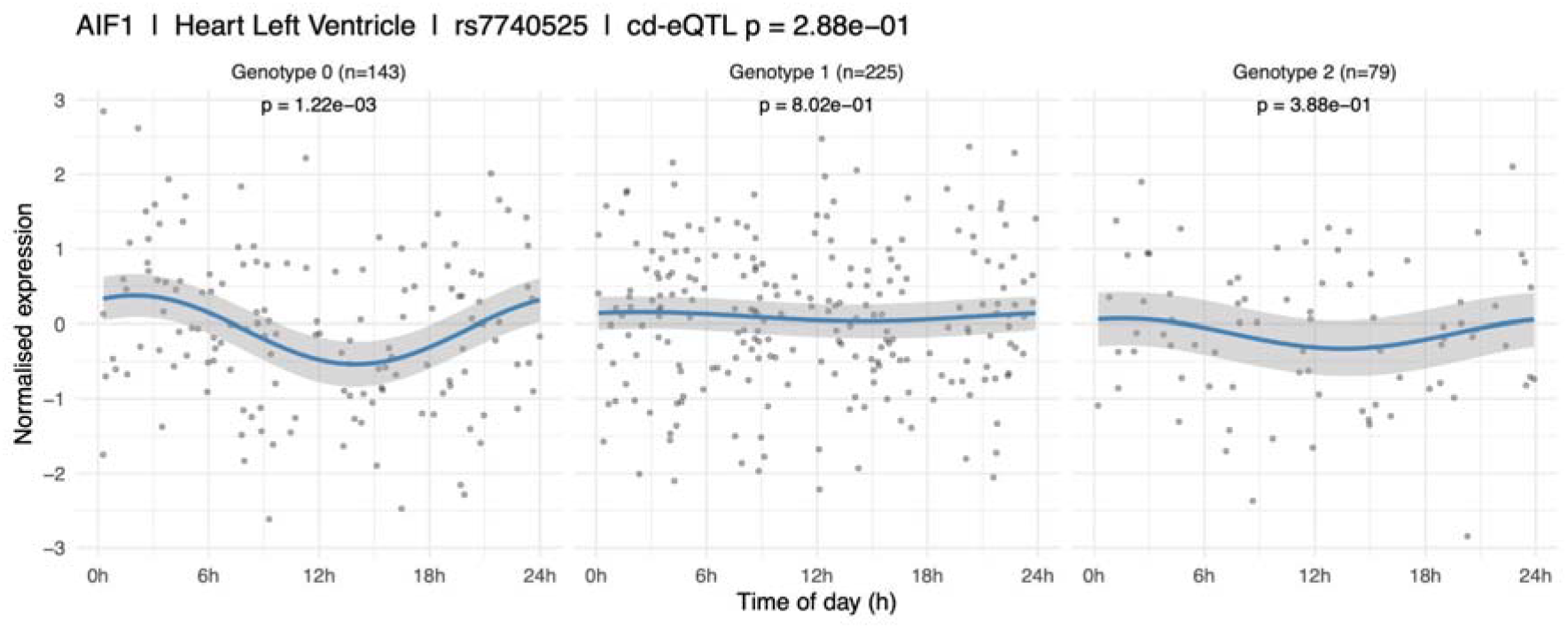
Genotype-stratified rhythmicity versus joint interaction model for *AIF1* in the heart left ventricle. Normalised *AIF1* expression in heart left ventricle plotted against time of day for each genotype group of rs7740525: homozygous reference (Genotype 0, n = 143), heterozygous (Genotype 1, n = 225) and homozygous alternate (Genotype 2, n = 79). Points are individual donors; the blue line is the fitted cosinor curve with its 95% confidence interval (grey band). The p-value above each panel is the within-group rhythmicity test, significant only in the homozygous reference group (p = 1.22 × 10^−3^) and non-significant in the heterozygous (p = 0.80) and homozygous alternate (p = 0.39). The joint genotype-by-time interaction model applied to the same data is not significant (p = 0.29).

## Notes

### Competing Interest Statement

The authors have declared no competing interest.

## References

1. Kuo, T. T., Wang, J. & Kay, S. A. Time matters: circadian genetics and the molecular logic of human health and disease. Nat. Rev. Genet. 27, 744–761 (2026).

2. Marcheva, B. et al. Circadian Clocks and Metabolism. Handb. Exp. Pharmacol. 127–155 (2013) doi:10.1007/978-3-642-25950-0_6.

3. Zhang, R., Lahens, N. F., Ballance, H. I., Hughes, M. E. & Hogenesch, J. B. A circadian gene expression atlas in mammals: implications for biology and medicine. Proc. Natl. Acad. Sci. U. S. A. 111, 16219–16224 (2014).

4. Guan, D. & Lazar, M. A. Circadian regulation of gene expression and metabolism in the liver. Semin. Liver Dis. 42, 113–121 (2022).

5. Talamanca, L., Gobet, C. & Naef, F. Sex-dimorphic and age-dependent organization of 24-hour gene expression rhythms in humans. Science 379, 478–483 (2023).

6. Fagiani, F. et al. Molecular regulations of circadian rhythm and implications for physiology and diseases. Signal Transduct. Target. Ther. 7, 41 (2022).

7. Yi, J. S., Díaz, N. M., D’Souza, S. & Buhr, E. D. The molecular clockwork of mammalian cells. Semin. Cell Dev. Biol. 126, 87–96 (2022).

8. Ueda, H. R. et al. System-level identification of transcriptional circuits underlying mammalian circadian clocks. Nat. Genet. 37, 187–192 (2005).

9. Yamamoto, T. et al. Transcriptional oscillation of canonical clock genes in mouse peripheral tissues. BMC Mol. Biol. 5, 18 (2004).

10. Hastings, M. H., Maywood, E. S. & Brancaccio, M. Generation of circadian rhythms in the suprachiasmatic nucleus. Nat. Rev. Neurosci. 19, 453–469 (2018).

11. Iiams, S. E., Skinner, N. J., Green, C. B. & Takahashi, J. S. Circadian Alignment Through Time-Restricted Feeding: Implications for Health and Longevity. Annu. Rev. Nutr. 46, 435–462 (2026).

12. Martin, R. A., Viggars, M. R. & Esser, K. A. Metabolism and exercise: the skeletal muscle clock takes centre stage. Nat. Rev. Endocrinol. 19, 272–284 (2023).

13. Jones, S. E. et al. Genome-wide association analyses of chronotype in 697,828 individuals provides insights into circadian rhythms. Nat. Commun. 10, 343 (2019).

14. Scheer, F. A. J. L., Hilton, M. F., Mantzoros, C. S. & Shea, S. A. Adverse metabolic and cardiovascular consequences of circadian misalignment. Proc. Natl. Acad. Sci. U. S. A. 106, 4453–4458 (2009).

15. Bass, J. & Takahashi, J. S. Circadian integration of metabolism and energetics. Science 330, 1349–1354 (2010).

16. Morris, C. J., Purvis, T. E., Hu, K. & Scheer, F. A. J. L. Circadian misalignment increases cardiovascular disease risk factors in humans. Proc. Natl. Acad. Sci. U. S. A. 113, E1402–1411 (2016).

17. Ector, C. et al. Circadian rhythm heterogeneity modulates drug response variations in neuroblastoma models. Cell Rep. 45, 116975 (2026).

18. Pariollaud, M. & Lamia, K. A. Cancer in the Fourth Dimension: What Is the Impact of Circadian Disruption? Cancer Discov. 10, 1455–1464 (2020).

19. Straif, K. et al. Carcinogenicity of shift-work, painting, and fire-fighting. Lancet Oncol. 8, 1065–1066 (2007).

20. Walker, W. H., Walton, J. C., DeVries, A. C. & Nelson, R. J. Circadian rhythm disruption and mental health. Transl. Psychiatry 10, 28 (2020).

21. Bunney, B. et al. Circadian dysregulation of clock genes: clues to rapid treatments in major depressive disorder. Mol. Psychiatry 20, 48–55 (2015).

22. THE GTEX CONSORTIUM. The GTEx Consortium atlas of genetic regulatory effects across human tissues. Science 369, 1318–1330 (2020).

23. Lonsdale, J. et al. The Genotype-Tissue Expression (GTEx) project. Nat. Genet. 45, 580–585 (2013).

24. Chen, Y., Liu, P., Sabo, A. & Guan, D. Human genetic variation determines 24-hour rhythmic gene expression and disease risk. Nat. Commun. 16, 4270 (2025).

25. Zhou, D. et al. Genetics-nutrition interactions control diurnal enhancer-promoter dynamics and liver lipid metabolism. Cell Metab. 37, 1961-1979.e7 (2025).

26. Taylor, J. & Tibshirani, R. J. Statistical learning and selective inference. Proc. Natl. Acad. Sci. 112, 7629–7634 (2015).

27. Pelikan, A., Herzel, H., Kramer, A. & Ananthasubramaniam, B. Venn diagram analysis overestimates the extent of circadian rhythm reprogramming. FEBS J. 289, 6605–6621 (2022).

28. Hutchison, A. L. et al. Improved statistical methods enable greater sensitivity in rhythm detection for genome-wide data. PLoS Comput. Biol. 11, e1004094 (2015).

29. Karczewski, K. J. et al. Pan-UK Biobank GWAS improves discovery, analysis of genetic architecture, and resolution into ancestry-enriched effects. 2024.03.13.24303864 Preprint at 10.1101/2024.03.13.24303864 (2024).

30. Balsalobre, A., Damiola, F. & Schibler, U. A serum shock induces circadian gene expression in mammalian tissue culture cells. Cell 93, 929–937 (1998).

31. Rieder, C. L. & Cole, R. W. Cold-shock and the Mammalian cell cycle. Cell Cycle 1, 169–175 (2002).

32. Davis, J. R. et al. An Efficient Multiple-Testing Adjustment for eQTL Studies that Accounts for Linkage Disequilibrium between Variants. Am. J. Hum. Genet. 98, 216– 224 (2016).

33. Weger, B. D. et al. Systematic analysis of differential rhythmic liver gene expression mediated by the circadian clock and feeding rhythms. Proc. Natl. Acad. Sci. 118, e2015803118 (2021).

34. Bingham, C., Arbogast, B., Guillaume, G. C., Lee, J. K. & Halberg, F. Inferential statistical methods for estimating and comparing cosinor parameters. Chronobiologia 9, 397–439 (1982).

35. Meireles-Filho, A. C. A., Bardet, A. F., Yáñez-Cuna, J. O., Stampfel, G. & Stark, A. cis-Regulatory Requirements for Tissue-Specific Programs of the Circadian Clock. Curr. Biol. 24, 1–10 (2014).

36. Qiu, P. et al. BMAL1 knockout macaque monkeys display reduced sleep and psychiatric disorders. Natl. Sci. Rev. 6, 87–100 (2019).

37. Daudali, H. et al. Genetic variation in circadian regulator gene BMAL1 in psychiatric, psychological and cardiometabolic traits: a trans-ancestry UK Biobank study. BMJ Ment. Health 27, e301267 (2024).

38. Cuddapah, V. A. et al. Rare variants in BMAL1 are associated with a neurodevelopmental syndrome. Proc. Natl. Acad. Sci. U. S. A. 122, e2427085122 (2025).

39. Edgar, R. S. et al. Cell autonomous regulation of herpes and influenza virus infection by the circadian clock. Proc. Natl. Acad. Sci. U. S. A. 113, 10085–10090 (2016).

40. Zhang, M. et al. A Translocation Pathway for Vesicle-Mediated Unconventional Protein Secretion. Cell 181, 637-652.e15 (2020).

41. Wang, Y. et al. TMED10-mediated unconventional secretion of IL-33 regulates intestinal epithelium differentiation and homeostasis. Cell Res. 34, 258–261 (2024).

42. Nagao, Y., Taguchi, A. & Ohta, Y. Circadian Rhythm Dysregulation in Inflammatory Bowel Disease: Mechanisms and Chronotherapeutic Approaches. Int. J. Mol. Sci. 26, 3724 (2025).

43. Weintraub, Y. et al. Clock Gene Disruption Is an Initial Manifestation of Inflammatory Bowel Diseases. Clin. Gastroenterol. Hepatol. Off. Clin. Pract. J. Am. Gastroenterol. Assoc. 18, 115-122.e1 (2020).

44. Wang, S. et al. REV-ERBα integrates colon clock with experimental colitis through regulation of NF-κB/NLRP3 axis. Nat. Commun. 9, 4246 (2018).

45. Preuss, F. et al. Adverse effects of chronic circadian desynchronization in animals in a “challenging” environment. Am. J. Physiol. - Regul. Integr. Comp. Physiol. 295, R2034–R2040 (2008).

46. Liu, J.-L. et al. Circadian Clock Disruption Suppresses PDL1+ Intraepithelial B Cells in Experimental Colitis and Colitis-Associated Colorectal Cancer. Cell. Mol. Gastroenterol. Hepatol. 12, 251–276 (2021).

47. Shimba, S. et al. Brain and muscle Arnt-like protein-1 (BMAL1), a component of the molecular clock, regulates adipogenesis. Proc. Natl. Acad. Sci. U. S. A. 102, 12071–12076 (2005).

48. Duez, H. & Staels, B. Circadian Disruption and the Risk of Developing Obesity. Curr. Obes. Rep. 14, 20 (2025).

49. Pocock, S. J., Ashby, D., Shaper, A. G., Walker, M. & Broughton, P. M. Diurnal variations in serum biochemical and haematological measurements. J. Clin. Pathol. 42, 172–179 (1989).

50. Jouffe, C. et al. Disruption of the circadian clock component BMAL1 elicits an endocrine adaption impacting on insulin sensitivity and liver disease. Proc. Natl. Acad. Sci. U. S. A. 119, e2200083119 (2022).

51. Kennaway, D. J., Varcoe, T. J., Voultsios, A. & Boden, M. J. Global loss of bmal1 expression alters adipose tissue hormones, gene expression and glucose metabolism. PloS One 8, e65255 (2013).

52. Shimba, S. et al. Deficient of a clock gene, brain and muscle Arnt-like protein-1 (BMAL1), induces dyslipidemia and ectopic fat formation. PloS One 6, e25231 (2011).

53. Toh, K. L. et al. An hPer2 phosphorylation site mutation in familial advanced sleep phase syndrome. Science 291, 1040–1043 (2001).

54. Zhang, L. et al. A PERIOD3 variant causes a circadian phenotype and is associated with a seasonal mood trait. Proc. Natl. Acad. Sci. U. S. A. 113, E1536–1544 (2016).

55. Archer, S. N. et al. A length polymorphism in the circadian clock gene Per3 is linked to delayed sleep phase syndrome and extreme diurnal preference. Sleep 26, 413–415 (2003).

56. Patke, A. et al. Mutation of the Human Circadian Clock Gene CRY1 in Familial Delayed Sleep Phase Disorder. Cell 169, 203-215.e13 (2017).

57. Hirano, A. et al. A Cryptochrome 2 mutation yields advanced sleep phase in humans. eLife 5, e16695 (2016).

58. Archer, S. N., Viola, A. U., Kyriakopoulou, V., von Schantz, M. & Dijk, D.-J. Inter-individual differences in habitual sleep timing and entrained phase of endogenous circadian rhythms of BMAL1, PER2 and PER3 mRNA in human leukocytes. Sleep 31, 608–617 (2008).

59. Hu, Y. et al. GWAS of 89,283 individuals identifies genetic variants associated with self-reporting of being a morning person. Nat. Commun. 7, 10448 (2016).

60. Jones, S. E. et al. Genome-wide association analyses of chronotype in 697,828 individuals provides insights into circadian rhythms. Nat. Commun. 10, 343 (2019).

61. Mermet, J. et al. Clock-dependent chromatin topology modulates circadian transcription and behavior. Genes Dev. 32, 347–358 (2018).

62. GTEx Consortium. Human genomics. The Genotype-Tissue Expression (GTEx) pilot analysis: multitissue gene regulation in humans. Science 348, 648–660 (2015).

63. Price, A. L. et al. Principal components analysis corrects for stratification in genome-wide association studies. Nat. Genet. 38, 904–909 (2006).

64. STAR: ultrafast universal RNA-seq aligner | Bioinformatics | Oxford Academic. https://academic.oup.com/bioinformatics/article/29/1/15/272537.

65. Robinson, M. D. & Oshlack, A. A scaling normalization method for differential expression analysis of RNA-seq data. Genome Biol. 11, R25 (2010).

66. Stegle, O., Parts, L., Piipari, M., Winn, J. & Durbin, R. Using probabilistic estimation of expression residuals (PEER) to obtain increased power and interpretability of gene expression analyses. Nat. Protoc. 7, 500–507 (2012).

67. Roenneberg, T. et al. Epidemiology of the human circadian clock. Sleep Med. Rev. 11, 429–438 (2007).

68. Zeileis, A. & Hothorn, T. Diagnostic Checking in Regression Relationships.

